# A formal relation between two disparate mathematical algorithms is ascertained from biological circuit analyses

**DOI:** 10.1101/2025.03.28.645962

**Authors:** Charles Liu, Elijah FW Bowen, Richard Granger

## Abstract

We simulate and formally analyze the emergent operations from the specific anatomical layout and physiological activation patterns of a particular local excitatory-inhibitory circuit architecture that occurs throughout superficial layers of cortex. The circuit carries out two effective procedures on its inputs, depending on the strength of its local feedback inhibitory cells. Both procedures can be formally characterized in terms of well-studied statistical operations: clustering, and component analyses, under high-feedback-inhibition and low-feedback-inhibition conditions, respectively. The detailed nature of these clustering and component procedures is studied in the context of extensive related literatures in statistics, machine learning, and computational neuroscience. The two operations (clustering and component analysis) have not previously been shown to contain deep connections, let alone to each be derivable from a single overarching algorithmic precursor. The identification of this deep formal mathematical connection, which arose from analysis of a detailed biological circuit, represents a rare instance of novel mathematical relations arising from biological analyses. ^1^

## 1 Key characteristics of superficial cortical circuitry produce specific algorithms

### 1.1 Bottom-up biological derivation of circuit components and assemblies

Most cortical regions incorporate superficial layer cell assemblies consisting of excitatory and inhibitory cells anatomically organized as in Figure 1. Evidence suggests that these local excitatory-inhibitory assemblies contain both sparse external cortico-cortical input axons, and dense reciprocal local connectivity between excitatory and local inhibitory cells (Pfeffer et al., 2013; Campagnola et al., 2022; Rudy et al., 2011; Tremblay et al., 2016; Douglas and Martin 2004; 2007).

**Figure 1:**
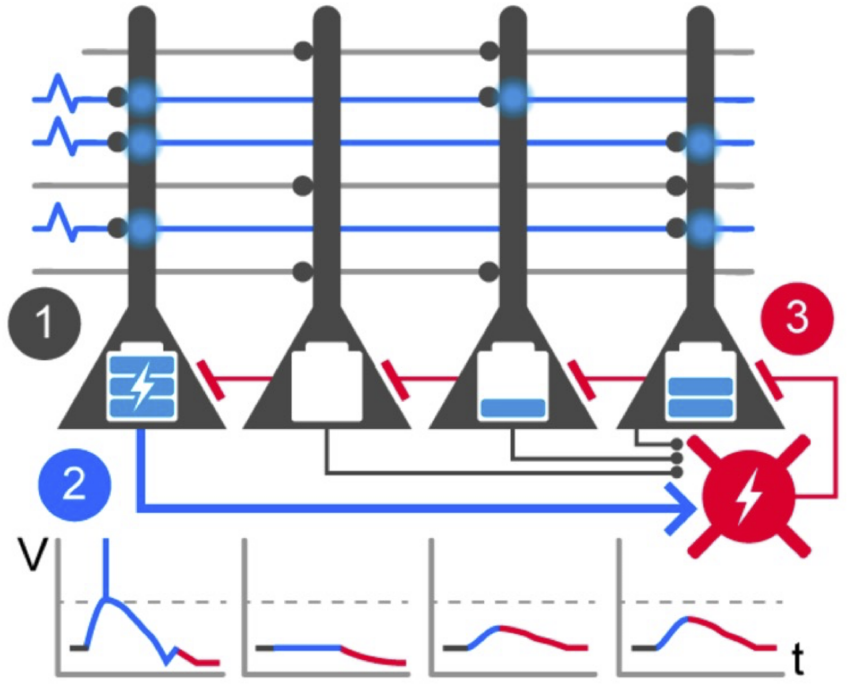
A cortical assembly composed of one inhibitory (red) and four excitatory (black) cells. Time-dependent voltage transients within the cells, shown as voltage (V) vs. time (t) graphs, build up over time as a function of the number of active input axons, and the number and weights of connected target dendritic synapses. The overall effect (see text) is that the most-activated excitatory cell (“1” in the figure) reaches its action potential threshold more rapidly than other cells; once any excitatory cell spikes, it activates (2) its local inhibitory cell (2), which in turn inhibits all local excitatory cells (3). Thus, only the most-activated excitatory cell(s) can respond (i.e., in a sense, the “winners” take all).

We forward a model of these assemblies that attempts to closely match both their anatomical layout and the nature and timing of their physiological interactions; we term these as local-feedback lateral inhibition circuits (L-FLIC).

These L-FLIC assemblies are connected such that (1) each glutamatergic input axon from other cortical cells, makes contact sparsely with a fixed number of excitatory target dendrites; and (2) each excitatory dendrite receives contacts from a separately fixed number of incoming axons (Felch 2008). Furthermore, (3) the excitatory cells densely contact local inhibitory cells (in a ratio of about 4:1) and (4) the inhibitory cells reciprocally make dense contacts with local excitatory cells (Abeles 1991; Fitzpatrick et al., 1987; Hendry et al., 1987).

Within a local L-FLIC assembly, when an activated excitatory neuron rises to spiking threshold, it activates its local inhibitory neuron, which then inhibits all other excitatory neurons within the radius of its local axonal arborization (which includes the excitatory cell that activated the inhibitory cell). Once an input pattern arrives spatially distributed across incoming axons, then all target excitatory neurons’ voltages rise, until the most rapidly rising cells attain their spiking threshold, at which point the emitted spikes activate the local inhibition, which inhibits all the local excitatory cells within its reach. This will halt the rise of voltage in the excitatory cells, so that any that have not yet spiked in response to the input, now will be unable to do so. Thus a very limited subset of target excitatory cells can successfully respond to an input.

The cells that do respond to a given input are determined by the speed of their voltage rise to their spiking threshold, which in turn is determined by the “fit” between the spatial pattern of incoming axon activity, and the distribution and conductivity of the sparsely distributed synapses on the dendrites of the target excitatory cells. Those cells with sufficiently conductive synapses that are connected to incoming axons that are active for this given input, are the cells that are likely to reach spiking threshold in response to this input.

### 1.2 Derivation of *competitive* circuit activity

The detailed unfolding of neuronal activity in these assemblies begins with an incoming set of spikes from axons arriving from some projecting cortical area. These in turn induce depolarization via input axons that contact dendritic synapses of neurons within the L-FLIC assemblies in the target cortical area. The total voltage over time in the target cells rises at a rate that is dependent on the quantity of input activation at a given target cells and the conductivity or weights of the receiving synapses. (These are depicted as voltage/time graphs in Figure 1).

Note also that the active synapses of the target cells will then become increased slightly in weight, due to synaptic plasticity. These synaptic increases then have an effect on any future encounters with this input pattern or with slight variants of the same input pattern. If synapses have increased their weights in response to a spatial input activation pattern *A*, then in response to slightly different pattern *A*^*′*^, many of the strengthened synapses will be activated (due to the similarity between *A* and *A*^*′*^), and that will generate a more rapid voltage rise in the target cell, making that cell more likely to reach its spiking threshold before being halted by feedback lateral inhibition. In other words, for any input that elicits some target cell spiking, variants of that input will also tend to elicit spikes from the same target cells, and synaptic plasticity further increases this probability of the same target cells responding to similar but somewhat-different inputs. As a result, similar inputs will increasingly come to elicit identical, not just similar, cortical responses.

Such an operation directly relates to clustering, a core unsupervised learning technique that partitions data into groups based on similarity. In clustering, data points are grouped into clusters such that those within the same cluster exhibit stronger similarities—such as higher inclination, shorter distances, or more alike features—compared to points in different clusters.

A set of information-processing operations is thus carried out directly from the anatomical organization and physiological time-sequence activity of a small assembly of well-studied excitatory and inhibitory cells. These assemblies are very distinct from standard artificial-neural-network (ANN) approaches to cell operation (e.g., Mao et al 2007; Kaski et al 1994; Amari 1977; Pinto et al., 2001; Oja 1989; Grossberg 1987; von der Malsburg 1973). Many of these ANN approaches can be characterized in terms of “winner take all” or “lateral inhibition”, and they bear some notable points of correspondence with the L-FLIC model, as well as several important differences, that will be discussed.

### 1.3 Derivation of *cooperative* circuit activity

Ascending systems, such as the norepinephrine system from the locus coeruleus, the acetylcholine system from the basal forebrain, and the dopamine systems from the ventral tegmental area (and the SNc), can inhibit cortical inhibitory cells (REFS). These systems are known to substantially participate in cortical activity via these modulatory inputs (Avery & Krichmar 2017) both to pyramidal cells and to interneurons (transiently inhibiting inhibition) (Jones 2003; Brown & McKenna 2015; Freund & Meskenaite 1992; Vazey et al., 2018; Kilavik et al., 2013; Kawaguschi et al., 1998; Salgado et al., 2011; Gulyas et al., 1996; Blasco-Ibanez et al., 1998; Gulyas et al., 1999).

In the L-FLIC circuit, what effects emerge in the presence of inhibited inhibition, i.e., suppression of the lateral inhibitory feedback cell in the L-FLIC assembly? Analysis in subsequent sections demonstrates and analyzes the activity under these circumstances. If feedback inhibition is weak or absent, then the initially-responding cells will still send signals to activate the local feedback inhibitory cell, but that inhibition now will be **damped or suppressed** by the ascending input and thus there will be little lateral feedback inhibition of local excitatory cells. In this case, multiple excitatory cells may succeed in reaching their spiking thresholds, in contrast with the relatively few (or single) excitatory responses when lateral inhibition is strong. As before, excitatory cells that do fire sufficiently strongly in response to these inputs are likely to experience induced synaptic change, i.e., the synapses receiving the activating inputs will become strengthened. Over trials, if many excitatory cells reach their spiking thresholds, and become further strengthened in these responses, what then is the information that this population of excitatory cells conveys? Intuitively, each cell can be considered as a synaptic vector, and synaptic change causes that vector to come to “point” more closely in the direction of the input activity vectors on which the responding cells were trained. Thus all of these excitatory cells will change direction as a function of the inputs they respond to. Which inputs do they respond to? In this case, almost all of them, since the local inhibitory cell has been weakened and cannot provide lateral inhibition that would select only certain responding excitatory cells. If the excitatory cells are thus being “trained” to respond to the population of inputs, these cells will tend to point in the general direction of the overall leading principal component of these inputs. This line of intuitive reasoning is subjected to formal treatment in subsequent sections. We show that the actual learned response can be considered as part of a general “family” of principal-component-like responses. The specific response is studied in detail, and is termed the “AIME” algorithm response.

### 1.4 Partial summary

In sum, i) there is a local excitatory-inhibitory circuit architecture that occurs widely throughout superficial layers of cortex; ii) this architecture is subject to ascending systems that may affect the strength of its inhibitory cells; iii) the circuit’s characteristic responses to structured (non-uniform) populations of inputs can be formally characterized, both in the high-feedback-inhibition and lower-feedback-inhibition cases. Both such responses turn out to succumb to analysis and show themselves to be interpretable in terms of well-studied statistical operations: clustering and component analysis. The specific nature of the clustering and component responses is derived and studied in detail, and the specific nature of the type of clustering, and the type of component analysis, that are performed are characterized.

Finally, we note that the two statistical operations of clustering and component analysis are not closely related in the statistics literature. In particular, there is no known generalized function that has clustering as one special case and component analysis as another special case. Yet we have shown that the specific biological L-FLIC architecture is exactly such a generalized function, with clustering and component analysis as two special cases of its operation. Thus a novel and unexpected unification has been found: a generalized architecture directly from neuroscience produces a set of operational steps whose special cases are two formal statistical algorithms that were not previously thought to be closely related to each other. The identification of a direct connection between these statistical methods is itself a novel finding; the derivation of that finding from brain circuit analysis may itself be of equal importance and interest.

## 2 Clustering within thalamocortical circuits

The cells thus far described in cortical superficial layers’ L-FLIC are combined with several additional layers of cells in neocortical circuitry (layers IV, V, and VI) and these are further incorporated into reciprocal thalamic structures, both thalamic “core” nuclei, and the overlaying inhibitory thalamic layer termed nucleus reticularis thalamus (NRT). We study how these added structures further interact with the superficial layer L-FLIC structure to perform an enlarged set of algorithmic operations. In particular, we show that for each of the identified fundamental L-FLIC operations, the additional layers enable the system to perform iteratively and hierarchically, i.e., successive iterations of clustering or component analysis.

The core thalamic nucleus first relays static input to the middle cortical layer (layer IV), which then transmits the information apically to the superficial layer (II-III), where the L-FLIC operations occur. While local lateral inhibition is present within the L-FLIC assemblies in the superficial layers, synaptic change over multiple trials eventually causes excitatory neurons to generate nearly identical responses to similar inputs from the middle layers, as already described.

The superficial layers then activate the deep layers (V-VI) in a topographic manner. Feedback from layer VI is then sent to both the thalamic reticular nucleus (NRT) and the core thalamus. The NRT inhibits portions of the core thalamic cells that correspond to just those cells already activated by the previous input. Due to the topographic organization, the inhibited portions also directly map back to the specific input regions that were just processed. This inhibition is long-lasting, persisting for hundreds of milliseconds (Huguenard and Prince, 1994; Cox et al., 1997; Zhang et al., 1997). As a result, whenever the same input is re-sensed, since it still is present in the environment, then only the uninhibited cells can be activated. That is, on successive sensory “samples” over time of a given input, first one set of cells is activated in thalamus (and then cortex), and then a quite different set of cells is activated on the second sampling, since the first set of target cells is now inhibited. We show that computationally, these sequential activities correspond to hierarchical clustering. We denote this process as lateral inhibition-hierarchical clustering (LI-HC).

LI-HC begins with an initial clustering and proceeds divisively through multiple learning cycles, each employing identical clustering modules connected by “input masking”. Given an input set *X* of size *p*-by-*n*, where each data point is a *p*-dimensional vector, the candidate winner set *W* contains sufficiently many *p*-dimensional weight vectors representing excitatory neurons. In each clustering module, weight vectors *w*_*i*_ are updated iteratively using the competitive learning rule:

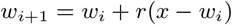

where *r* is the learning rate. Over iterations, the weight vectors gradually align with the input vector *x*. As this process is repeated for all data points, weight vectors associated with spatially similar inputs converge toward their mean, forming well-separated clusters.

After initial clustering, due to the long-lasting inhibition from GABAergic NRT overlaying the core thalamic cells, the next sample of the same static input will contain only the secondary portion of input activity that was not shared among the members of the established cluster. Now the previous winners and their corresponding input portions become dormant. This means that the input is partially “masked” for further learning, and the previously learned feature is “forgotten” by presently active excitatory neurons within L-FLIC.

They are now only likely to be activated by the remaining unmasked information. This masking initiates a transition across cycles in the learning process. Following the same learning rule, the “active remainder” of the input actions will choose a different set of excitatory target neurons compared to whatever was selected in the previous cycle. After training, the new neural weight vector will point from the parent cluster’s (determined in the previous cycle) centroid toward the unique features present in this specific input instance. This process is known as sub-clustering, a step following the initial clustering in LI-HC. The training process is simplified in the Algorithm 1. This will be shown to be an elaboration and variant of a mechanism studied by Rodriguez et al (2004).

### Algorithm 1 LI-HC

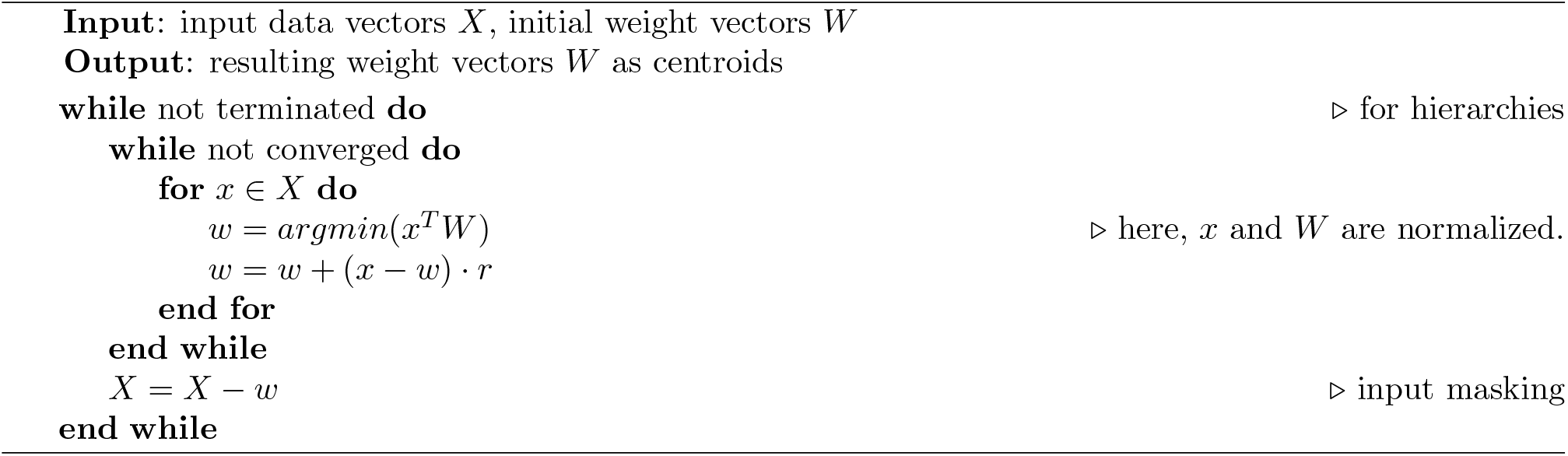

Geometrically, “input masking” is essentially vector subtraction: the input subtracts the winners from all previous cycles they have won, with the head of the previous winning vector becoming the tail of the current winner. In other words, input masking is equivalent to origin shifting, which concludes the previous cycle of clustering as well as initiates the next round of hierarchical clustering. This learning process ends when there is no further separable data or upon a pre-defined termination criterion. Ultimately, all the selected and updated winners become the final centroids for each hierarchy of clusters in the process. Figure 2 illustrates a sample result of LI-HC ^2^.

**Figure 2:**
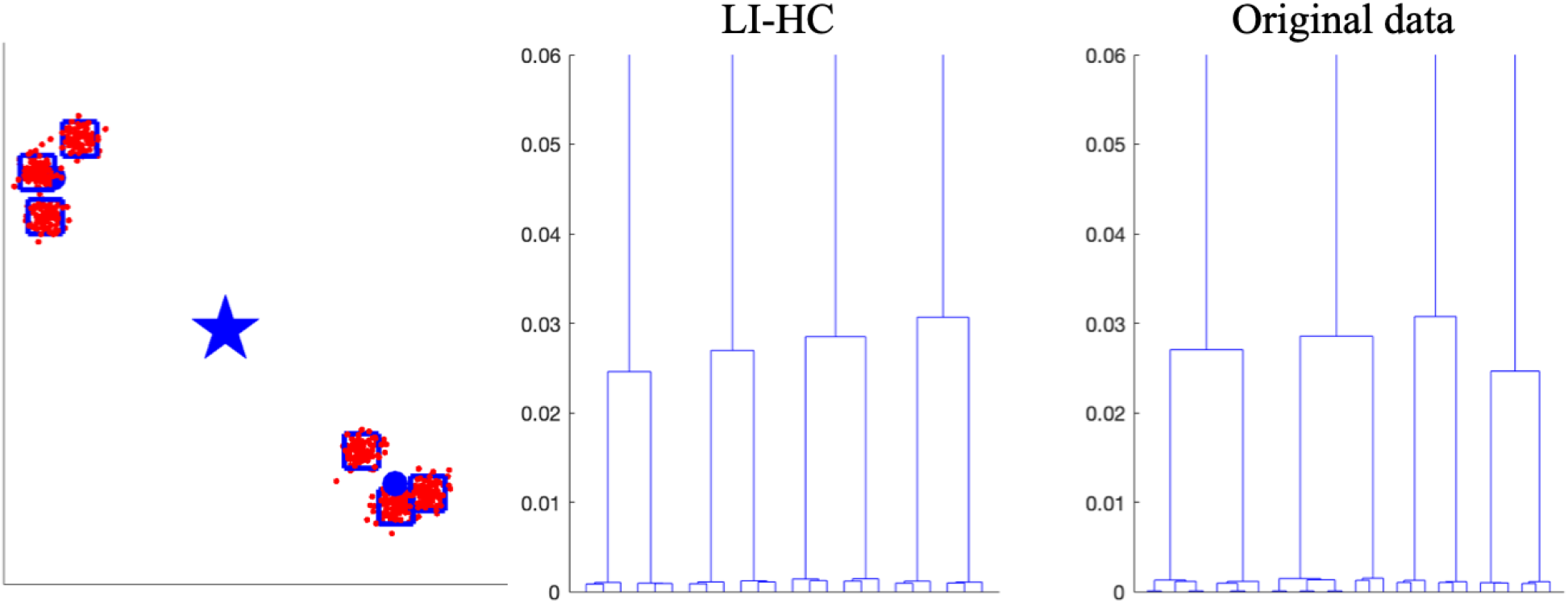
Demonstration of LI-HC. The leftmost figure presents the graphical outcome of LI-HC applied to a dataset containing two main clusters, each further subdivided into three sub-clusters. In this visualization, stars, dots (the upper left dot is partially overlayed under the other symbols), and hollow squares represent the learned weight vectors, serving as centroids at different hierarchical levels. The middle and rightmost figures display dendrograms constructed from the learned centroids by LI-HC and from the original data, respectively. Notably, the hierarchical cut-offs observed along the y-axes exhibit a high degree of similarity, indicating that the learned centroids effectively capture the hierarchical categories of the original data.

## 3 AIME algorithm

### 3.1 Significance

Recall that a specific small adjustment to the L-FLIC circuit: making the inhibition in the superficial layer of L-FLIC weak enough to be negligible – the local feedback circuits then perform PCA-like feature extraction. We here formalize the details of this feature extraction, introducing the algorithm termed “AIME”, Antipodal Iterative Mean Estimation. We show that the AIME algorithm has significant characteristics both in biological and computational terms.

- Biologically, AIME enhances the diversity of activities within local feedback circuits, due to different inhibitory patterns. In the context of inhibited inhibition, neurons respond to exterior stimuli without engaging in competitive behavior, and each neuron gains independent and identical access to these inputs. While operating within the same anatomical architecture, this transformation from the typical competitive excitatory-inhibitory patterns represents a fundamental change in physiological dynamics.
- Computationally, AIME expands the range of feature extraction techniques in data science and machine learning, providing a more generalized approach to capturing meaningful components. Principal Component Analysis (PCA) is a widely used data analysis technique that simplifies complex datasets by identifying key patterns and reducing dimensionality (Jolliffe & Cadima, 2016). Traditional PCA methods provide analytical and highly faithful results, with the first principal component consistently associated with the largest eigenvalue. This approach effectively captures the direction of maximum variance in the data and only provides a single set of axes for uncovering underlying structures and reducing dimensionality. However, when data is evenly distributed across multiple axes, several directions may explain the variance comparably well. In such cases, it is more meaningful to consider a family of lines through the data, rather than just a single set of axes. AIME is proposed to address this issue by producing a more comprehensive representation of the data’s underlying spreading. In sum, AIME is proposed as a generalized variant of the process of component analysis.

### 3.2 Stage 1: Identification of initial AIME components

AIME trains the neural weight vectors in two stages, each reflecting a different type of feedback loops. The first stage, inspired by the cooperative activities in L-FLIC, adjusts the weight vectors based on the most prominent distribution of the input data. The second stage, modeled after additional layers of neocortical circuitry, operates hierarchically to obtain a full set of representations of data dispersion.

In stage 1 (or the first learning cycle), AIME learns the original input data until convergence, with the main focus on identifying the direction of its maximum spread. During the process, each weight vector is updated toward the spatial axes defined by the data points, with the sign() function ensuring it forms acute angles with these axes (the right angles can be ignored due to the orthogonality). At the end of stage 1, the algorithm produces a set of *learned weight vectors*, which capture the primary “components” of the original input. It is important to note that what truly matters for illustrating data spread are the lines/axes (regardless of the directions and magnitudes) on which these learned weight vectors lie, and we refer to these lines as the *AIME components*. It is worth noting that in stage 1, we allow multiple AIME components to be obtained. In this paper, “lines” and “AIME components” may be used interchangeably. The learning rule of AIME is highly simplified and similar to that in LI-HC, but it exhibits distinct properties:

1. AIME prohibits normalization of the input data. Compressing the input data to the unit circle destroys its original spread pattern, thus voiding the significance of meaningful components.
2. The conventional “winner-take-all” effects fail to hold due to the absence of neural competition. This change allows each neural weight vector a substantial degree of independence for training and updates. Without competitive dynamics, neurons no longer appear inclined to local categories within the input data. Instead, each neuron is capable of capturing information from the entire input set, resulting in a more holistic representation of the data.
3. The learning focus shifts from “values” to “axes”. Specifically, the weight vectors can learn not only the exact values of the data points but also the bi-directional axes where the data points lie. This expanded learning capacity enables the algorithm to encode more spatial information about the input data rather than achieving categorization.

### 3.3 Stage 2: Identification of remaining AIME components

In stage 2, AIME performs hierarchical computations to extract additional meaningful components that are orthogonal to the leading components identified in stage 1. This process is analogous to determining successive principal components (PCs) based on the first PC. In contrast to the single-cycle stage 1, stage 2 consists of multiple learning cycles. For example, given a *p*-dimensional input set, stage 1 extracts the AIME components corresponding to the first dimension, while stage 2 performs the remaining *p*−1 learning cycles to extract subsequent components.

Specifically, at the beginning of *c*-th cycle (1 *< c* ≤ *p*), the algorithm performs an “input masking” step to enable continuous learning. This step involves projecting the current data onto a hyperspace orthogonal to the AIME components identified in the last cycle. By doing so, the leading AIME components of the current cycle are secondary to those obtained in the previous cycle. In other words, each subsequent learning cycle aims to identify the direction of maximum spread of the *transformed* input data. This iterative adjustment ensures hierarchical learning progression and comprehensive identification of all AIME components. Figure 3 illustrates this process by showing how the data is transformed to calculate deeper AIME components. The complete AIME algorithm with a single neural weight vector is provided in Algorithm 2.

**Figure 3:**
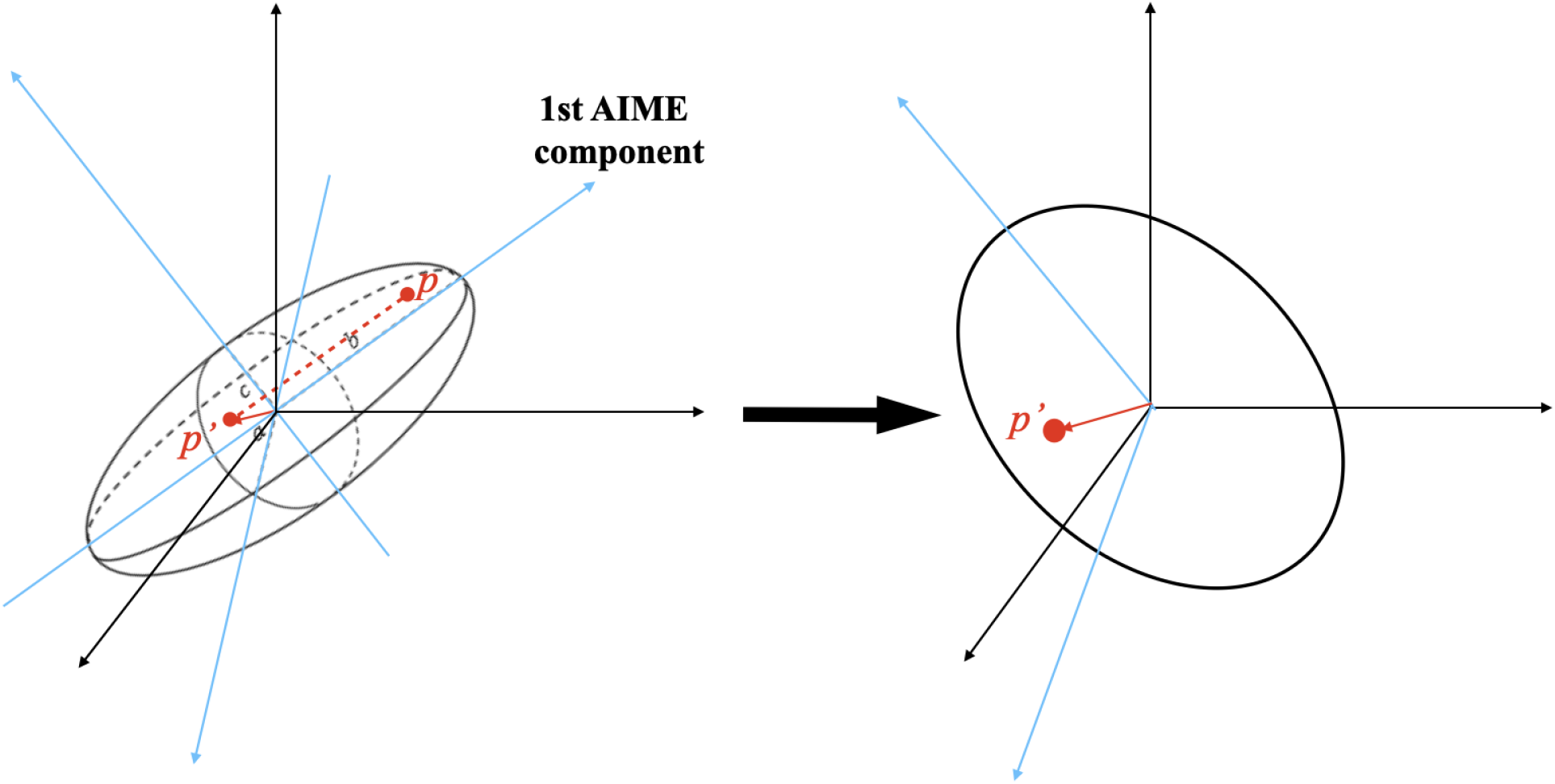
A geometric illustration of “input masking” using the first AIME component. The left subfigure shows data points distributed within an ellipsoid, where the polar radius *b* represents the first AIME component identified in the initial learning cycle. To compute the next AIME component, data points are projected onto a plane orthogonal to *b*, corresponding to the ellipsoid’s cross-section through the origin. For a data point *p* and the first AIME component *v*, the projection is calculated as 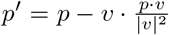. The right subfigure demonstrates that this transformation reshapes the data into a 2D “circular region” within the 3D space, facilitating further analysis.

#### Algorithm 2 AIME with a single weight vector

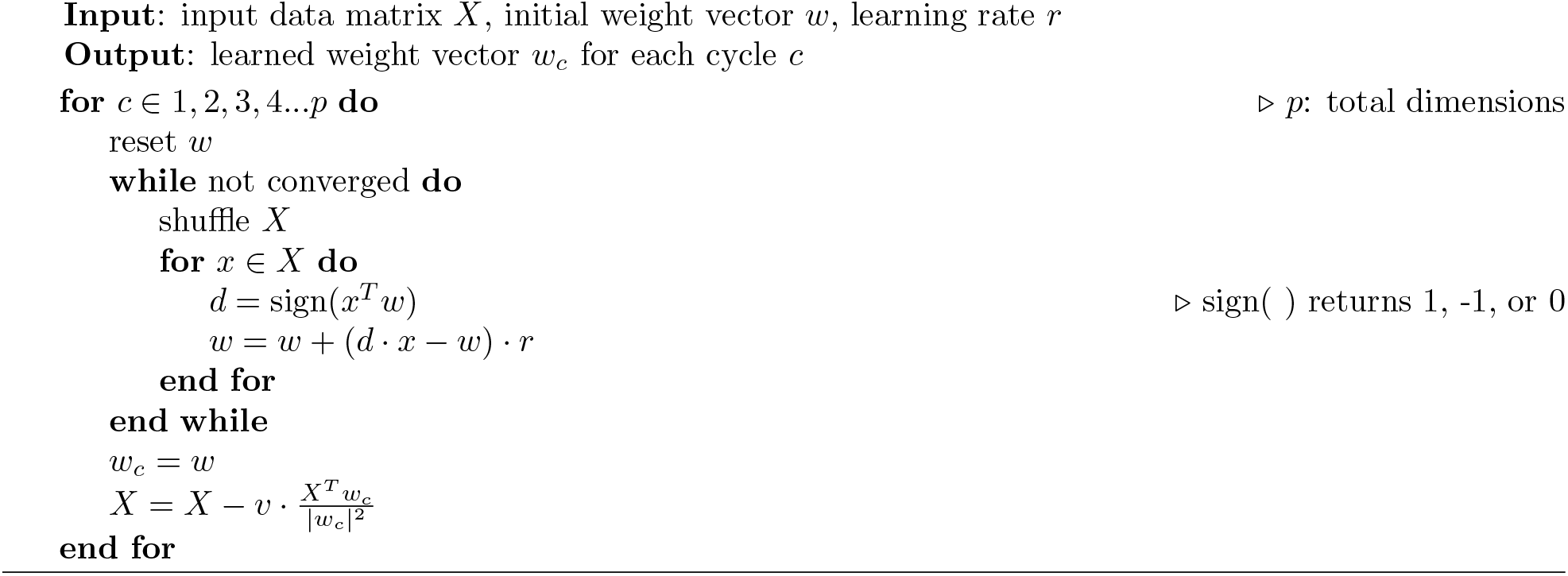

## 4 AIME mathematics

In this section, we carefully justify the AIME algorithm both with formal treatment and with empirical data-driven analyses. We provide both extensive simulations to illustrate the behavior of the algorithm, and a sketch of a proof that allows the derivation of the algorithm from sound mathematical foundations. A successful implementation of the algorithm should align with the following assumptions.

1. We assume that the input set is mean-centered and no data point is exactly located at the origin.
2. The size of the neural weight vectors and the input data should be sufficiently large (this is explained further).
3. It is mandatory to guarantee that the input data is not perfectly uniform along all axes of the hyperplane. Equivalently, the variances should not be identically explained by all principal components - the data is assumed to have some structure in the input space. (If the data does not contain structure to be dicovered, all results will simply be degenerate.)

### 4.2 Definitions and initial proof derivations

Consider the input set *X*_*n×p*_ (for simplicity, we write it as *X*), which contains *n* data points {*x*_1_, *x*_2_, …, *x*_*n*_}, each with *p* dimensions. The **Antipodal Augmentation** of *X* is the union of *X* and its negation, −*X*, and is written as

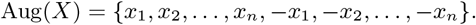

Geometrically, Aug(*X*) ∈ ℝ^*p*^ represents the bi-directional expansion of *X* in the coordinate, where each point is mirrored across the origin (see Figure 4:(a)). In an *p*-dimensional Euclidean space ℝ^*p*^, a hyperplane *H* passing through the origin is defined as:

**Figure 4:**
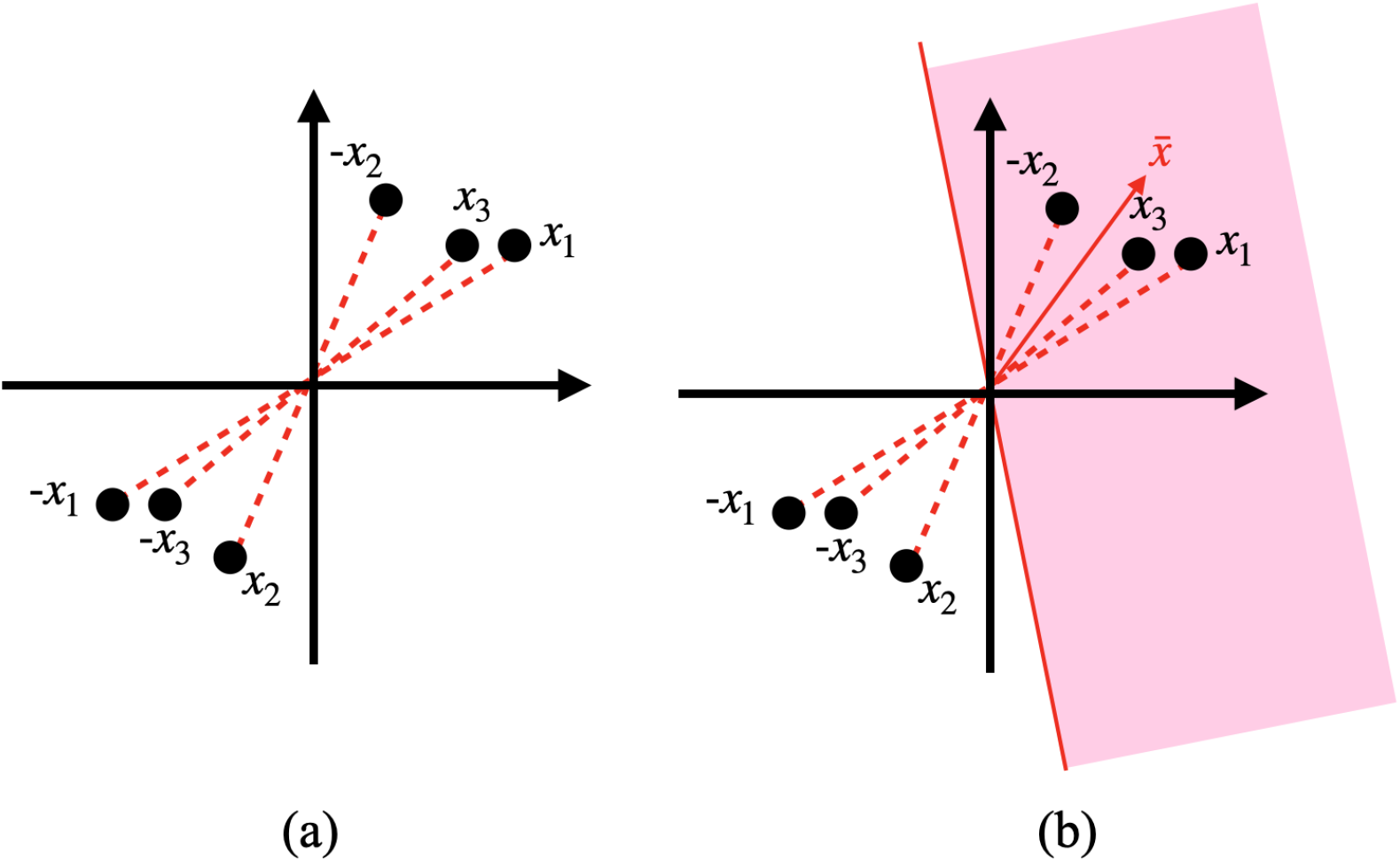
Antipodal augmentation of a sample input set *X*_3*×*2_ and an acute partition. **(a)** shows the antipodal augmentation of *X*, where *X* = {*x*_1_, *x*_2_, *x*_3_} and Aug(*X*) = {*x*_1_, *x*_2_, *x*_3_, −*x*_1_, −*x*_2_, −*x*_3_}. In **(b)**, the pink panel is an acute partition of Aug(*X*) with the corresponding 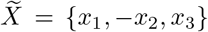. Their mean vector, 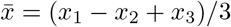, is acute to each of these vectors.

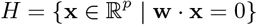

where **w** ≠ **0** is a normal vector. This hyperplane bipartitions ℝ^*p*^ into two open half-spaces:

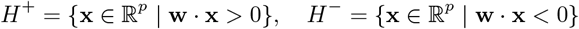

In Aug(*X*), each such half-space is defined as a **partition** of Aug(*X*). In each partition, either *x*_*i*_ or −*x*_*i*_ (∀*i* ∈ {1, 2, 3, …, *n}*) is included. We denote the set of vectors in a partition of Aug(*X*) as 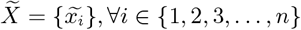, where each 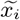 is either *x*_*i*_ or −*x*_*i*_. The data points on the partitions formed by the same hyperplane are symmetric according to the origin. For the discussion of existence and uniqueness, we consider partitions that contain the same set of vectors as equivalent. Therefore, there are a total of 2^*n*^ distinct partitions of Aug(*X*). Additionally, we define an **iteration** as a complete training epoch when AIME possesses the entire set of input data points. During each iteration, the sign() function determines the set of 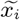. Therefore, the AIME algorithm is essentially an iterative search for an optimal partition, terminating once a specified condition is met.

A partition of Aug(*X*) is called an **acute partition** (denoted as *ap*) if its associated vectors *x*-_*i*_ form acute angles with their mean (see Figure 4:(b)). These vectors are referred to as the **acute-partition vectors**, and the mean of these vectors is called the **acute-partition mean**. For simplicity, we say that the acute-partition vectors are “acute to” their acute-partition mean. In Aug(*X*), each vector *x*_*i*_ has an associated antipodal pair, consisting of *x*_*i*_ and −*x*_*i*_. These two vectors lie on the same straight line and the collection of all such lines, formed by antipodal pairs, intersect at the origin. Each of these lines is referred to as the **line of** *x*_*i*_ or equivalently, the **line of** −*x*_*i*_. We now propose a heuristic algorithm to find an acute partition of an arbitrarily given input set *X* (satisfying the assumptions stated earlier) in Algorithm 3.

#### Algorithm 3 Acute Partition Finding Heuristic

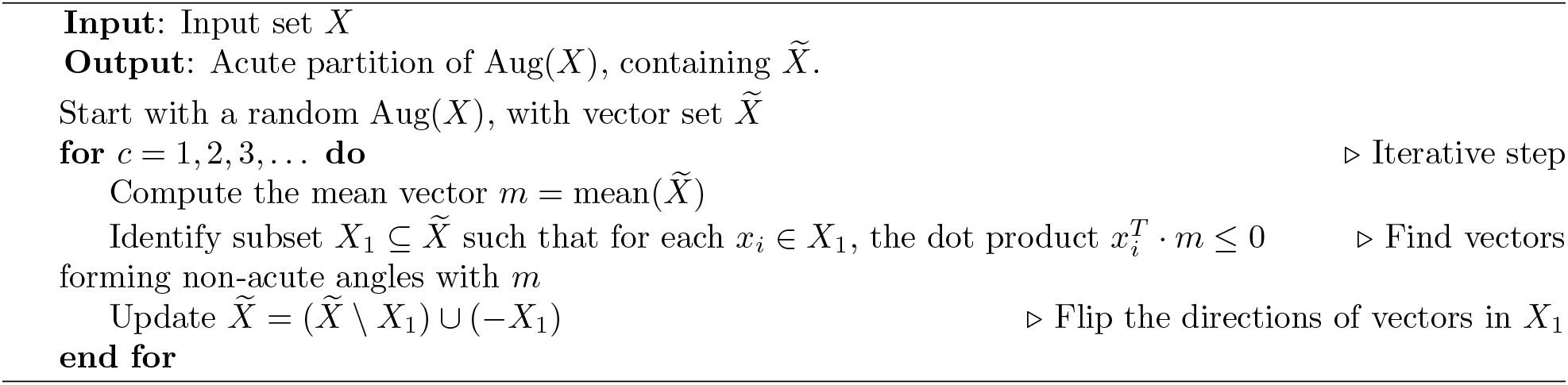

**Lemma** (Existence of an acute partition). For any *X*_*n×p*_ with *n >* 1 and *p >* 0, after *c* iterations, the probability that an acute partition exists in the antipodal augmentation of *X* approaches 1 as *c* increases. In other words, in probability, there exists at least one acute partition on the antipodal augmentation of any real-valued input set satisfying the assumptions. Mathematically,

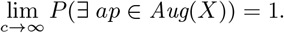

*Proof*. We use the Bayes’ theorem to illustrate the reasoning of the Lemma, which is equivalent to validating Algorithm 3. In the heuristic algorithm, the replacement step

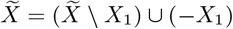

updates 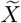 by replacing a subset of its elements with their negations. Intuitively, this step works by gradually eliminating non-acute effects in the partition. If an acute partition has not been found yet, the *i*th replacement can result in two possible outcomes:

1. Conditioned on the previous results, the present replacement does not yield an acute partition. Denote such an outcome as *N*_*i*_.
2. Conditioned on the previous results, the present replacement successfully creates an acute partition. Denote such an outcome as *F*_*i*_.

As the algorithm proceeds, after sufficiently many iterations *c*, the probability of not finding an acute partition after all replacements is given by the product of conditional probabilities *P* (*N*_1_ ∩ *N*_2_ ∩ … ∩ *N*_*c*_) = *P* (*N*_1_) · *P* (*N*_2_|*N*_1_) · *P* (*N*_3_|*N*_1_ ∩ *N*_2_) · · · · · *P* (*N*_*c*_|*N*_1_ ∩ *N*_2_ ∩ … ∩ *N*_*c*−1_). Due to the arbitrariness of input data, each probability lies in [0, 1], and not all of them equal 1. Consequently, the probability of finding at least one acute partition is:

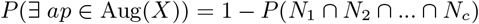

As the number of iterations *c* increases, the product of these probabilities becomes negligibly small, leading to:

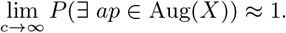

Figure 5 shows the simulated percentage of acute angles as the algorithm progresses, with a sample artificial dataset that has been deliberately designed to be evenly distributed to imitate the worst scenarios.

**Figure 5:**
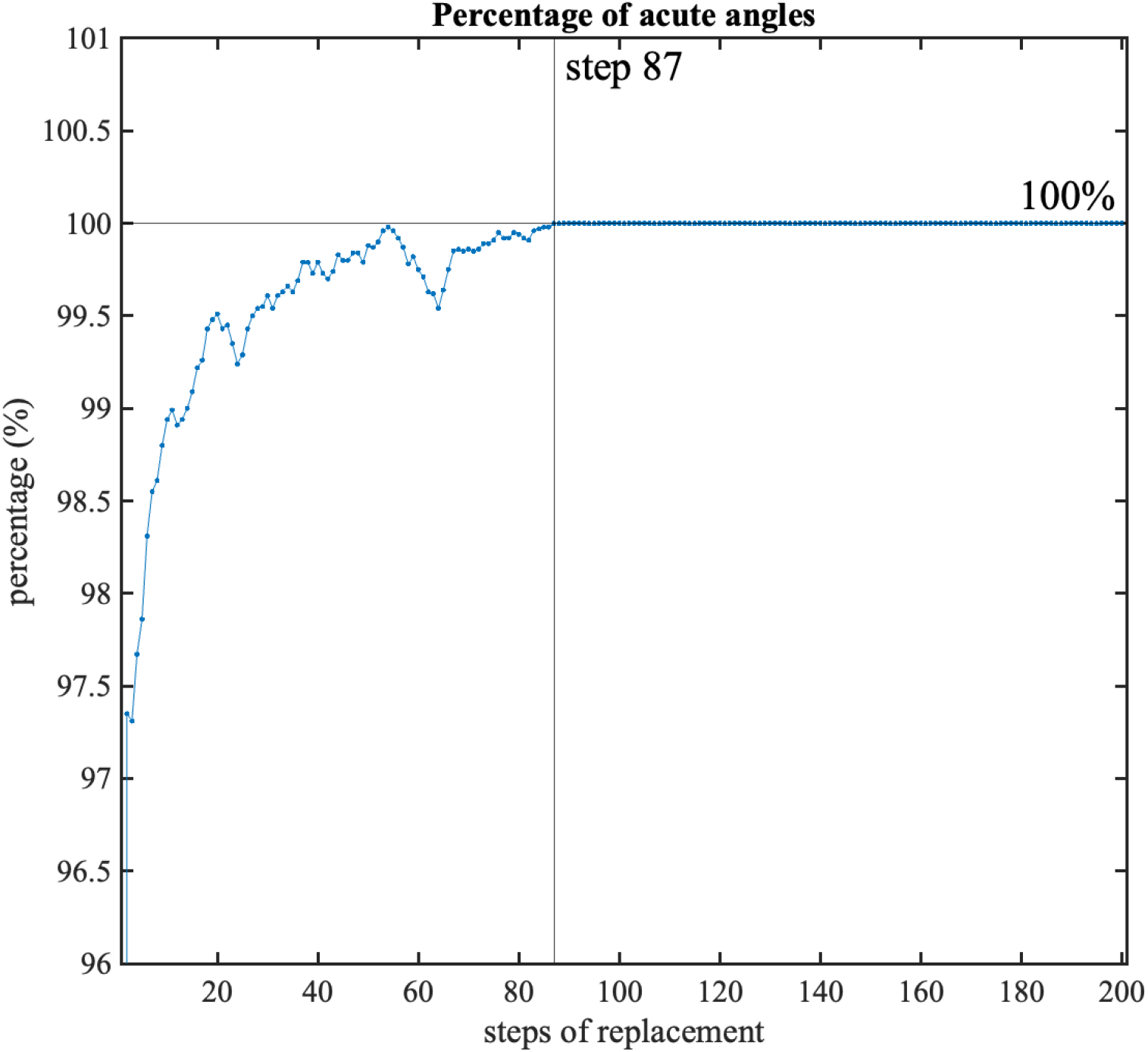
Simulation of the percentage of acute angles. It illustrates the process of finding an acute partition via Algorithm 3. Each “step of replacement” determines a new set of vectors 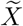. Each point on the curve represents the percentage of acute angles among the intersection angles between 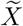 and its mean vector. The acute partition is found when this percentage constantly stays at 100%.

### 4.2 Single-neuron convergence

For simplicity, we first discuss the scenario where only a single neural weight vector is involved. With the AIME algorithm, the following statement justifies the convergence.

**Proposition 1** (Weak Law of Convergence). Under the AIME algorithm, with a sufficiently large number of iterations denoted by *c* on the input set *X*, a randomly initialized weight vector *w*_0_ (whose subscript indicates the index of iterations) will move towards the mean of the vectors it is acute to, and converge at the mean of an acute partition within Aug(*X*), where subtle oscillations do not induce changes in 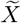 on the partition.

*Proof*. We prove the statement by strong induction.

1. **Basis step:** This step is the start of convergence. In the first iteration, the weight vector *w*_1_ selects a partition of the antipodal augmentation where the angles between *w*_1_ and the selected vectors are acute. The right angles can be ignored because the sign() function in the algorithm returns zero for orthogonal vectors, making no changes in that updating step. Recall that each iteration consists of *n* updating steps, where each step represents a weight update using an individual data point. These updates can cause the angle between the updated weight vector and its previously interacted 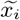 to shift from acute to obtuse (or vice versa), leading to a redefinition of the current partition. However, with a minimal learning rate *r*, each updating step is infinitesimal, where the set of 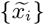 is assumed to stay fixed within the first iteration. With the fixed target vectors, the process becomes identical to the clustering module of LI-HC, where the weight vector moves toward the mean of the selected vectors in this partition. Hence, the basis case is valid.
2. **Inductive hypothesis:** Assume the statement holds for all integers *m* = 1, 2, 3, …, *k*, where *k* ≥ 2. Specifically, for any *m*-th iteration, *w*_*m*_ moves towards the mean of the vectors it is acute to, and this convergence is achieved at the *k*-th step when *k* is sufficiently large.
3. **Inductive step:** At the *k*-th step, by the inductive hypothesis, the weight vector *w*_*k*_ has approximately reached the mean of the vectors in its acute partition due to the assumed convergence. Subtle oscillations around the convergence point do not alter the partition or the corresponding vectors, as stated in the problem. Now, consider the (*k* + 1)-th iteration. Since the learning rate *r* → 0 as *k* → ∞, the angle adjustments in each updating step during this iteration are negligibly small, as demonstrated with the law of sines (illustrated in Figure 6:(a)). With further training over multiple updates, the cumulative effect of these small adjustments is further averaged out, ensuring that *w*_*k*+1_ remains asymptotically stationary. At this point, *w*_*k*+1_ remains acute to all vectors in the current partition. By induction, the statement holds for all integers *m >* 0. This completes the proof of the weak law of convergence. □

**Figure 6:**
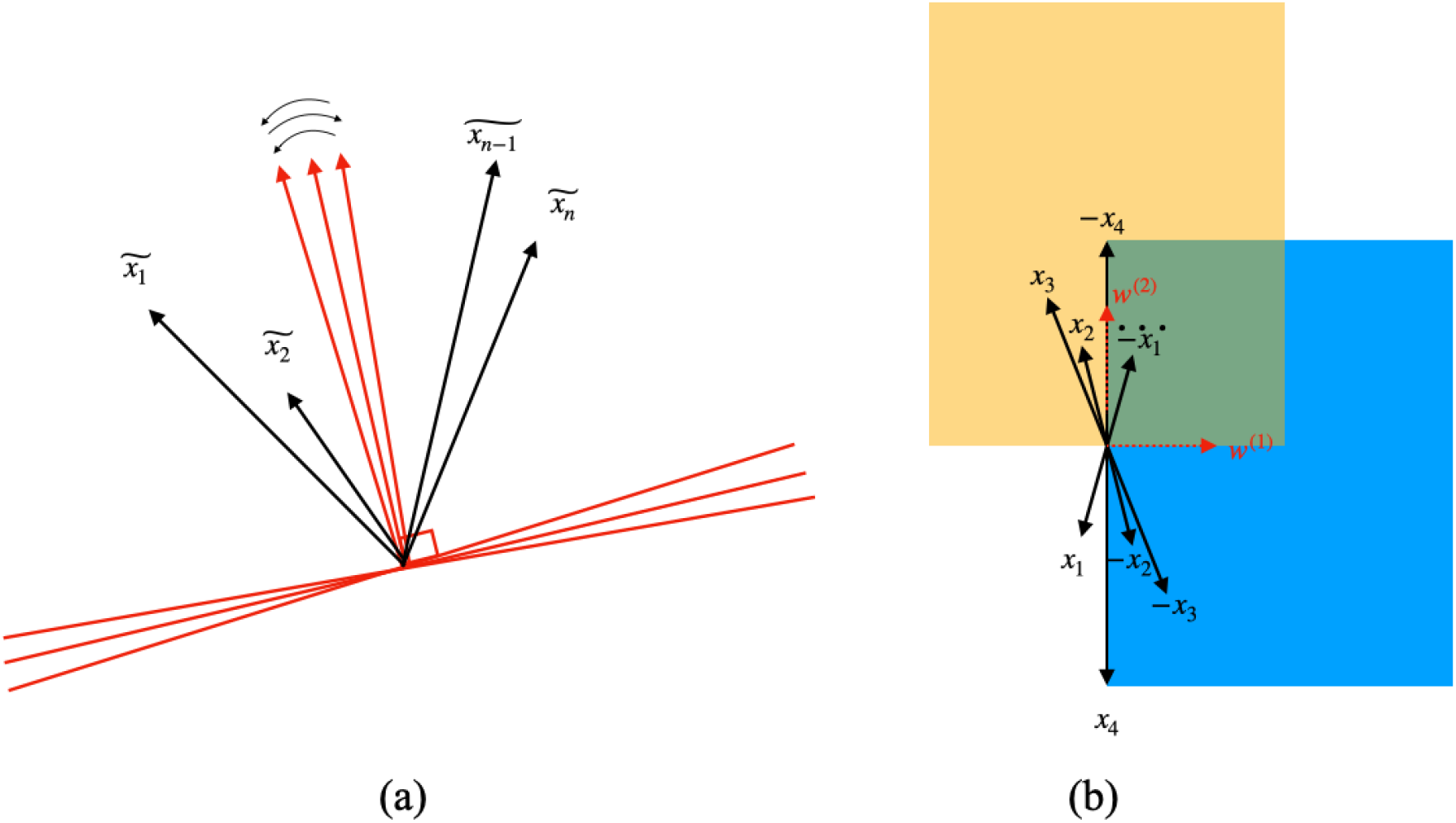
Laws of convergence. **(a)** shows the terminating phase upon convergence. Additional iterative steps (sufficiently small due to a small *r*) would trigger the weight vector to oscillate around the point of convergence, but will not include a new set of vectors. The data points of the current 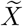 are fixed. **(b)** The convergence is more likely to be achieved at the yellow panel (at *w*^(2)^) rather than the blue one (at *w*^(1)^). This is because, in the blue panel, a small change caused by a single iterative step can more easily introduce new vectors, thus breaking the presumed convergence. Here, the yellow panel is a stronger acute partition than the blue one.

Although acute partitions exist, their uniqueness is not guaranteed. This raises an important question: with potentially multiple acute partitions, to which specific acute-partition mean will the weight vector ultimately converge? In other words, which acute partition ensures that subtle oscillations do not cause changes in the associated vectors? Indeed, the specific convergence point is highly data-driven and influenced by the spatial arrangement of the datasets. **Qualitatively**, the weight vectors tend to converge towards the acute partitions where the data points of 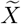 are more tightly concentrated around their mean.

We term such a partition with the densest concentration of vectors around the mean as the “strongest acute partition”, where the mean vector (denoted as strongest acute partition mean, or **“SAPM”**) maximizes the degree of strength of such vector-mean concentration. We propose the following qualitative statement to locate the convergence point more robustly given a single weight vector.

**Proposition 2** (Strong law of convergence). Under the AIME algorithm, with a sufficiently large number of iterations denoted by *c* on the input set *X*, a randomly initialized weight vector *w*_0_ will converge towards the SAPM.

*Proof*. We prove this statement by contradiction. Suppose that convergence occurs at a weaker acute partition, where part of 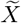 forms significantly larger angles from the mean. In such a weaker acute partition, the partition includes either −*x*_*i*_ or *x*_*i*_ for all *i* ∈ {1, 2, 3, …, *n}*. As a result, this acute partition divides the region where the vectors are densely distributed. Due to this dense distribution, small oscillations in the weight vector during further iterations are highly likely to cause the selection of new sets of vectors. This, in turn, results in the creation of a new 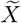, thus a change of partition. Such a shift contradicts the stationary phase of convergence, where the weight vector and its associated partition are expected to remain stable. Figure 6:(b) further illustrates this contradiction: a small perturbation in the blue panel (a weaker partition) may cause the inclusion of *x*_2_ and *x*_3_ while excluding −*x*_2_ and −*x*_3_, leading the resultant weight vector to shift to the yellow panel. This implies that the weaker acute partition cannot be the terminating partition. Therefore, by contradiction, the weight vector must converge to the mean of the strongest acute partition, where small oscillations do not alter the partition. This completes the proof. So far, we have qualitatively demonstrated the convergence of a single weight vector: it terminates at the SAPM.

### 4.3 Multi-neuron convergence

We now extend single-neuron findings to multiple neurons. In the strongest acute partition, the vectors exhibit the highest inclination toward their mean, known as the SAPM. The strength of this convergence can be measured in various ways, with one effective metric assessing how well the mean vector captures the variability within the partition. This aligns closely with PCA, particularly the explained variance of the vectors relative to the mean vector. Given this shared metric, we propose that the strongest acute partition is more likely to yield the largest projected variances, just like what PCA does.

However, if the projected variances along different axes are comparable, multiple SAPM solutions may exist. In contrast, the leading principal component of *X* is always unique. Therefore, we conclude that the line formed by the leading principal component, *p*_1_, belongs to the family of lines formed by SAPM (these lines are indeed AIME components), denoted as *S*, i.e., *p*_1_ ∈ *S*. When a group of neural weight vectors is given, each one follows the convergence rule elaborated above and terminates at an SAPM. There are two possible results.

1. First, if the input data is well structured such that the SAPM uniquely exists, all the learned weight vectors will approximately aggregate along a single AIME component, which approximately aligns with the leading principal component (PC) of the input data. In other words, *S* = {*p*_1_}. We verify the statement by contradiction: if it does not align with the leading PC, it indicates that the leading PC does not explain the biggest variance. This is in conflict with the definition of PC.
2. Second, when the data is approximately evenly distributed across the space, there are potentially tied findings of SAPM. In this case, the learned weight vectors appear on multiple AIME components and one of them is parallel to the leading PC. i.e., *p*_1_ ∈ *S*, where *S* ≠ {*p*_1_}.

With increasing dimensions and size, stronger non-linearity, and known distribution patterns of the input data, the likelihood of noise introduction increases, which potentially compromises the accountability of the single axis. These observations significantly challenge the uniqueness of the “strongest acute partition” and may cause the points of convergence to be located on multiple lines. This concern underscores the necessity for an abundant supply of initial weight vectors. Building on this, our algorithm, AIME, serves two functions: (1) AIME primarily aims to estimate the most accurate true principal components (PCs), as it consistently derives lines parallel to the true PCs. (2) However, when the true PCs fail to fully capture the variability in the data, with a sufficient number of weight vectors, AIME exhibits the capability to capture additional supplemental components with comparable strength. In other words, despite different directions, the variances explained along these axes show a remarkable similarity. The requirement for an expanded set of weight vectors is consistent with the biological hypothesis that more than one excitatory neurons get activated with a weaker excitation-inhibition pattern.

### 4.4 AIME, PCA, and Linear Regression

In summary, the core mathematical principles of “AIME” are reflected in its name:

- **A**ntipodal: The algorithm operates on the augmented dataset Aug(*X*).
- **I**terative: Learning occurs through an iterative process.
- **M**ean: The weight vector converges to the mean of the strongest acute partition.
- **E**stimation: AIME always produces decent estimates of the true principal components, regardless of the data distribution.

Our mathematical analysis and data-driven simulations show that at least one set of AIME components closely aligns with standard PCs, suggesting that the eligible AIME components share key properties of PCs. Linear regression, another widely used statistical technique, estimates linear relationships between data features (Freeman David A, 2009). In some cases, the leading PC and regression line may appear similar, leading to the mistaken assumption that they are equivalent. However, this is incorrect for several reasons. Since AIME components mirror PCs, the key comparison among these three algorithms—AIME, PCA, and regression—ultimately reduces to the fundamental distinction between PCA and regression. For the sake of this comparison, we consider PCA and AIME to be equivalent.

Firstly, when data is not centered at the origin, PCA/AIME and linear regression clearly show diverse outcomes.

- Principal components and AIME components pass through the origin due to mean-centering, while the regression line typically has a non-zero intercept as it minimizes errors in the dependent variable.
- The linear regression will always pass the mean of data due to its very calculation process, while the leading PC may not point to the data mean (Gallagher, O’Sullivan, & Palacios, 2020). This applies similarly to the AIME algorithm. Specifically, if the input data is not centered at the origin, the AIME line passes the input mean if and only if either Aug(*X*) = *X* or Aug(*X*) = −*X*.

Secondly, even with origin-centered data, PCA/AIME and linear regression remain mathematically distinct. Linear regression finds the best-fit line by minimizing the sum of squared vertical errors between predicted and actual values (Yan Xin, 2009). In contrast, PCA/AIME identifies the direction that maximizes data variance, equivalently minimizing the orthogonal distances from data points to the component line.

In short, PCA and AIME are closely related, while both differ significantly from linear regression.

## 5 Simulation methods

### 5.1 Data preparation

#### Input sets

To accurately implement the AIME algorithm and clarify its relationship with the true principal components, we run simulations using two different types of datasets: Iris data with a sharp spatial structure and synthetic data with a more even distribution. The Iris dataset (Fisher, 1988) stands as a classic and illuminating dataset, widely employed across a variety of machine learning tasks, including classification, clustering, and PCA. The Iris data comprises 150 samples from three distinct species of Iris (categorical: Iris setosa, Iris virginica, and Iris versicolor), with each sample characterized by four features (numerical), namely, the length and width of sepals and petals. This paper takes the four numerical features as the variables to ensure the continuity of the input data. The left sub-figure in Figure 7 shows the original Iris data alongside the first three principal components in a 3-dimensional space. Simultaneously, Table 1 lists the explained variances for each principal component, highlighting the sharp decreases observed between consecutive PCs of the Iris data.

**Table 1:** Explained variances of both the Iris data and the synthetic data. Two sub-tables display the first seven principal components of the data (if applicable) both individually and cumulatively. PCs of the Iris data show steep gradients, with PC1 explaining 92.2% of the variance and the following PCs explaining variances three times greater than the next PC. Whereas in the synthetic data, the explained variances progressively decrease through principal components. Notably, the synthetic data’s fourth and fifth PCs explain nearly identical variance percentages, indicating that the data spreads more evenly along these two axes.

| data | PC1 | PC2 | PC3 | PC4 | PC5 | PC6 | PC7 | ... |
| --- | --- | --- | --- | --- | --- | --- | --- | --- |
| Iris data | 92.2% | 5.4% | 1.8% | 0.5% | N/A | N/A | N/A | N/A |
| Synthetic data | 31.5% | 12.9% | 8.2% | 6.8% | 6.1% | 4.5% | 2.9% | ... |

| data | PC1 | PC2 | PC3 | PC4 | PC5 | PC6 | PC7 | ... |
| --- | --- | --- | --- | --- | --- | --- | --- | --- |
| Iris data | 92.2% | 97.6% | 99.5% | 100% | N/A | N/A | N/A | N/A |
| Synthetic data | 31.5% | 44.4% | 52.6% | 59.4% | 65.5% | 70.0% | 72.9% | ... |

**Figure 7:**
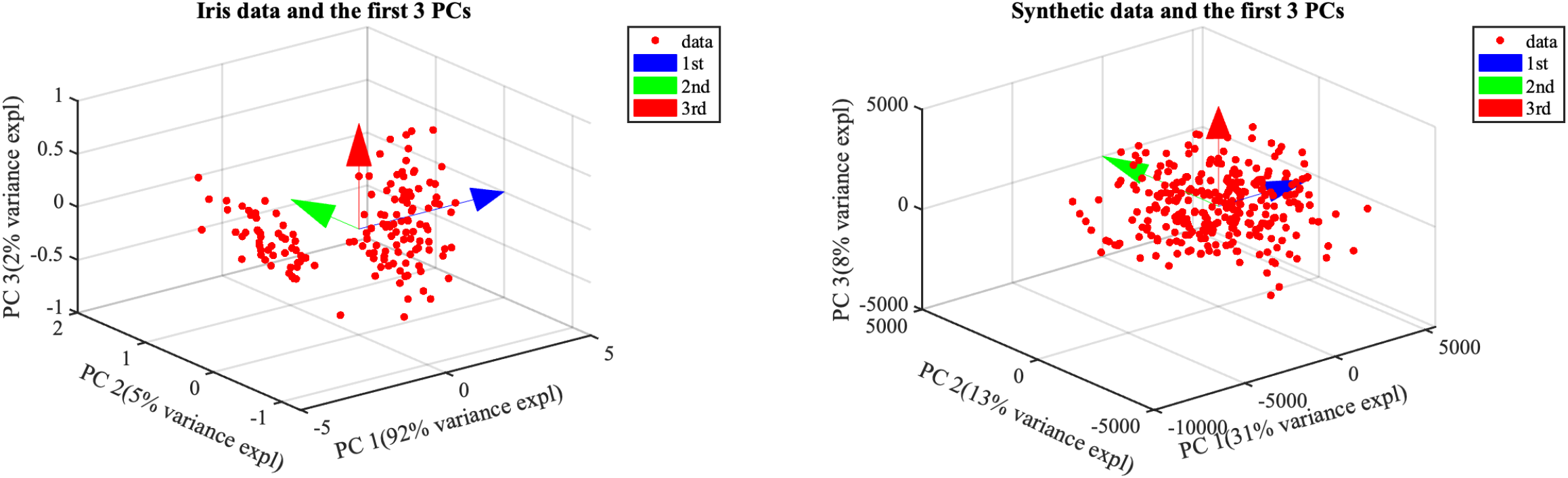
Input data sets with their first three true principal components. **Left:** Iris data, 4-dimensional with 150 data points. **Right:** synthetic data, 50-dimensional with 300 data points.

Secondly, we apply AIME to a synthetic input dataset with 300 instances and 50 numerical features (Schork J., n.d.). The outstanding difference is that the Iris data is well-structured, with a sharp gradient in explained variances of PCs. On the contrary, synthetic data presents an evenly distributed pattern, as shown in the right sub-figure of Figure 7 and evidenced by the slowly decreasing variances presented in Table 1.

#### Neural weights initiation

Excitatory neurons in the L-FLIC are sparsely connected in synaptic networks. How is such sparsity mathematically modeled? Felch and Granger (2008) point out that the sparse synaptic networks should follow a hypergeometric distribution to minimize a range of computational costs. Therefore, this paper initializes the synaptic vectors by strategically placing zeros based on a hypergeometric distribution. While the AIME algorithm is versatile and capable of functioning with a variety of randomly initialized weight vectors, the incorporation of hypergeometric patterns significantly enriches its biological interpretability.

### 5.2 AIME components determination

After each learning cycle, the obtained AIME components relate to the leading principal component in two ways. (1) When the input data is organized in a way that a specific direction can explicitly explain the maximum variance in the data, all learned weight vectors tend to appear along the same line. In this case, the AIME component is uniquely determined and the leading PC aligns with the line. Imagine this like a spotlight shining directly on the most important direction of variability in the data. Each learned weight vector, representing the strongest concentration of the data, corresponds directly to this spotlighted direction and its counter-direction. (2) When the input data lacks such a clear structure and is more evenly distributed in space, the learned weight vectors may be aggregated on a family of lines (i.e., multiple AIME components are obtained from a single learning cycle). All of these lines yield similar explained variances of the input data. The leading PC is approximately parallel to one of these lines. This scenario reflects a more complex interplay of different axes explaining the data’s variability.

With such complex scenarios within a single learning cycle and in combination with hierarchical operations, the determination of the full spectrum of AIME components involves two key steps:

- **Step 1: line finding within each cycle**. The raw outcomes directly from the training of each cycle are the learned weight vectors, and these vectors aggregate into groups. In theory, we should expect that these groups are symmetric at the origin, and the line connecting two “antipodal groups” should (approximately) pass through the origin. To determine these groups, K-Means clustering (or other types of user-defined clustering methods) is applied to precisely categorize the resulting vectors, where *K* is specified after subjective observation. This approach helps locate the aggregated lines (i.e., AIME components) and clarify their organization.
- **Step 2: input masking with inheritance**. The current input data is then projected onto the hyperplane (passing the origin) perpendicular to each AIME component determined in Step 1 for “input masking.” Step 1 may involve obtaining multiple lines that explain comparable variances, and the input data is then transformed based on each of these lines. The child AIME components are directly inherited from their parent components and are meaningful only within that context. They cannot be compared or analyzed across different parent components, as there are no cross-relationships between them. For example, given the parent AIME components *line*_*A*_ and *line*_*B*_, the child AIME components derived from input data masked upon *line*_*A*_ have no meaningful comparison with *line*_*B*_ and its children.

### 5.2 Measurements

The validation of AIME involves two measurements from both geometric and numerical perspectives. Firstly, the **correlation** is used to evaluate the geometric relationship between the true principal components and the AIME components. We focus solely on the parallelism of these lines, irrespective of their pointing direction. Therefore, this paper retains only the correlations’ absolute values, ranging from 0 to 1. If there are multiple AIME components, we show the results of each line.

In conjunction with correlation, the **cumulative explained variance** (in %) provides a numerical examination of the dispersion of data along the lines (true PCs and AIME components). When dealing with high-dimensional data, this paper simplifies stage 2 of the algorithm by focusing only on the first few components that capture most of the total variance. This approach avoids an exhaustive discussion of all dimensions and results in rescaled percentages of explained variance.

## 6 Simulation results

### Summary

The results from the Iris dataset and the synthetic dataset reveal distinct patterns, clearly demonstrating the following:

1. Validity of AIME – AIME is an effective algorithm for estimating true PCs as well as extracting other meaningful components if needed.
2. Well-structured Iris Data – The Iris dataset has a clear structure, with contrasting concentrations along different dimensions. As a result, AIME produces lines closely matching with true PCs.
3. Noisy Synthetic Data – The synthetic dataset represents a noisier scenario, with data more evenly distributed and lacking clear spatial patterns. Consequently, AIME generates a family of components, one of which aligns with the true PCs.

### Diagram

Figure 8 outlines the resulting lines (i.e., AIME components) produced by Iris and the synthetic data. Iris data’s dispersion is tight and the learned weight vectors roughly form a single AIME component after each cycle. With the synthetic data, which is more uniformly distributed across different axes, we obtain multiple AIME components. The diagram highlights input masking with inheritance.

**Figure 8:**
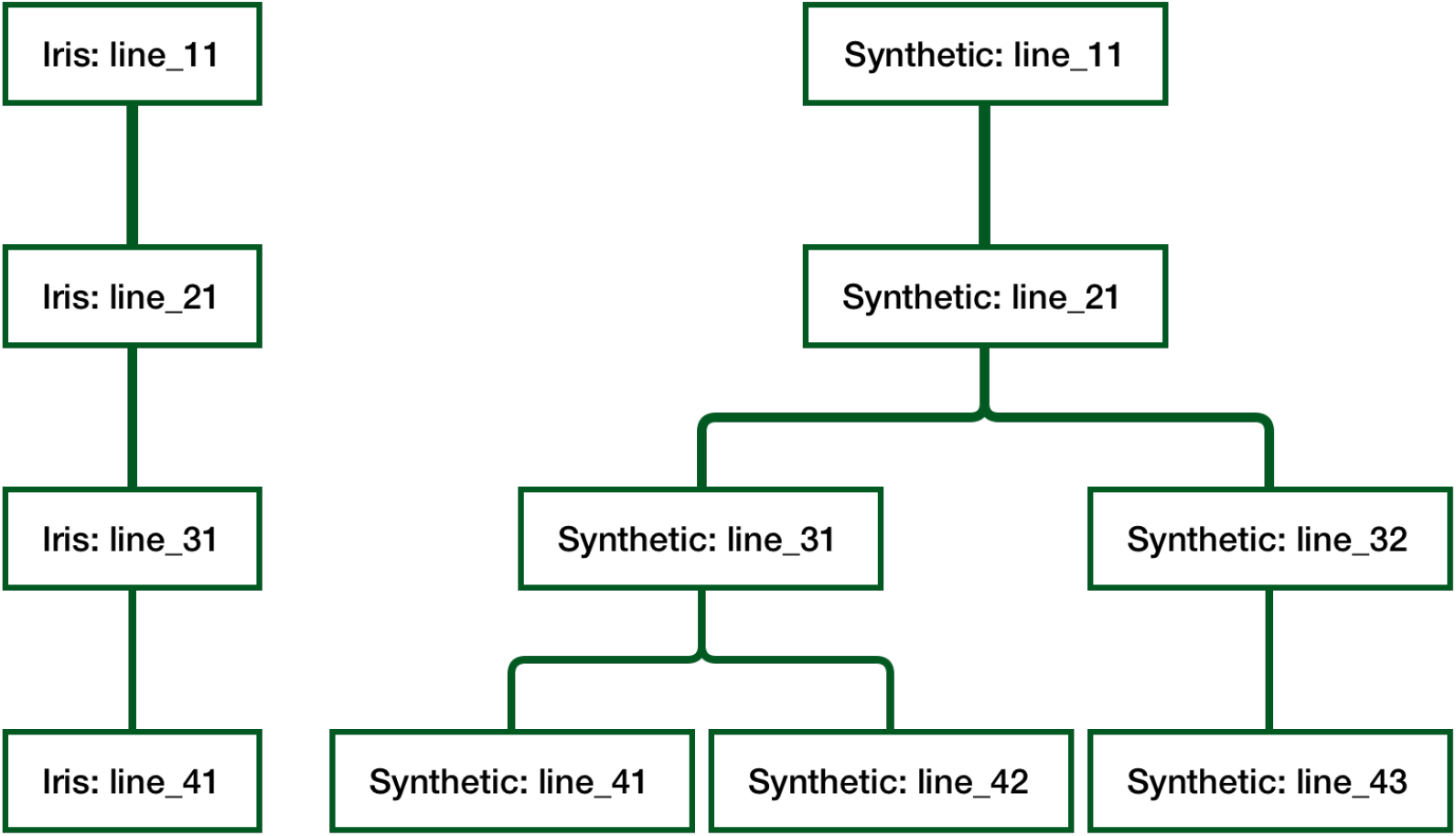
The learning diagram depicts the progression of both datasets using the AIME algorithm across multiple cycles, with hierarchical branches representing inherited input masking. The left-hand side (LHS) diagram illustrates the learning process using the Iris dataset, where each cycle produces a single AIME component showing a straightforward linear learning trajectory. In contrast, the right-hand side (RHS) diagram portrays the learning process with synthetic data, resulting in a tree-like structure due to the emergence of multiple distinct AIME components in cycles 3 and 4. In the end, three sets of AIME components are obtained: {*line*_11_, *line*_21_, *line*_31_, *line*_41_, *line*_11_, *line*_21_, *line*_31_, *line*_42_}, and {*line*_11_, *line*_21_, *line*_32_, *line*_43_}. This complication in the RHS diagram highlights the diverse representations of data dispersion across deeper dimensions.

### Graphical outcomes

Figure 9 and 10 show the learned weight vectors (aggregated in groups) and their formed lines with both datasets. These graphical results are consistent with the diagrams.

**Figure 9:**
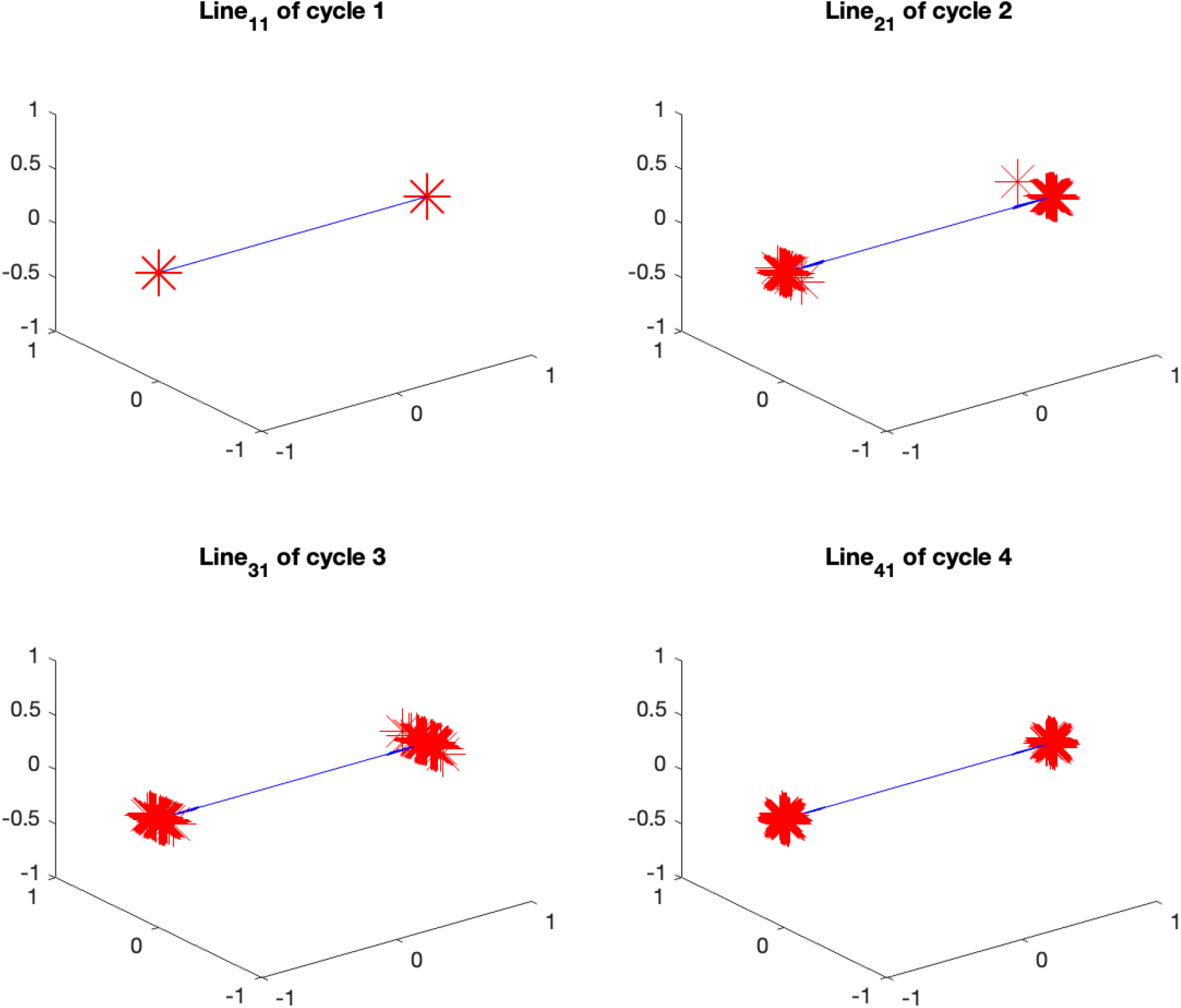
Learned weight vectors forming a unique AIME component with Iris data after each cycle. After each learning cycle, the learned weight vectors roughly appear along a single AIME component (in dark blue) spanning two directions. This line, determined by K-Means clustering with *K* = 2, aligns with the leading principal components of the current input data (either original or masked).

**Figure 10:**
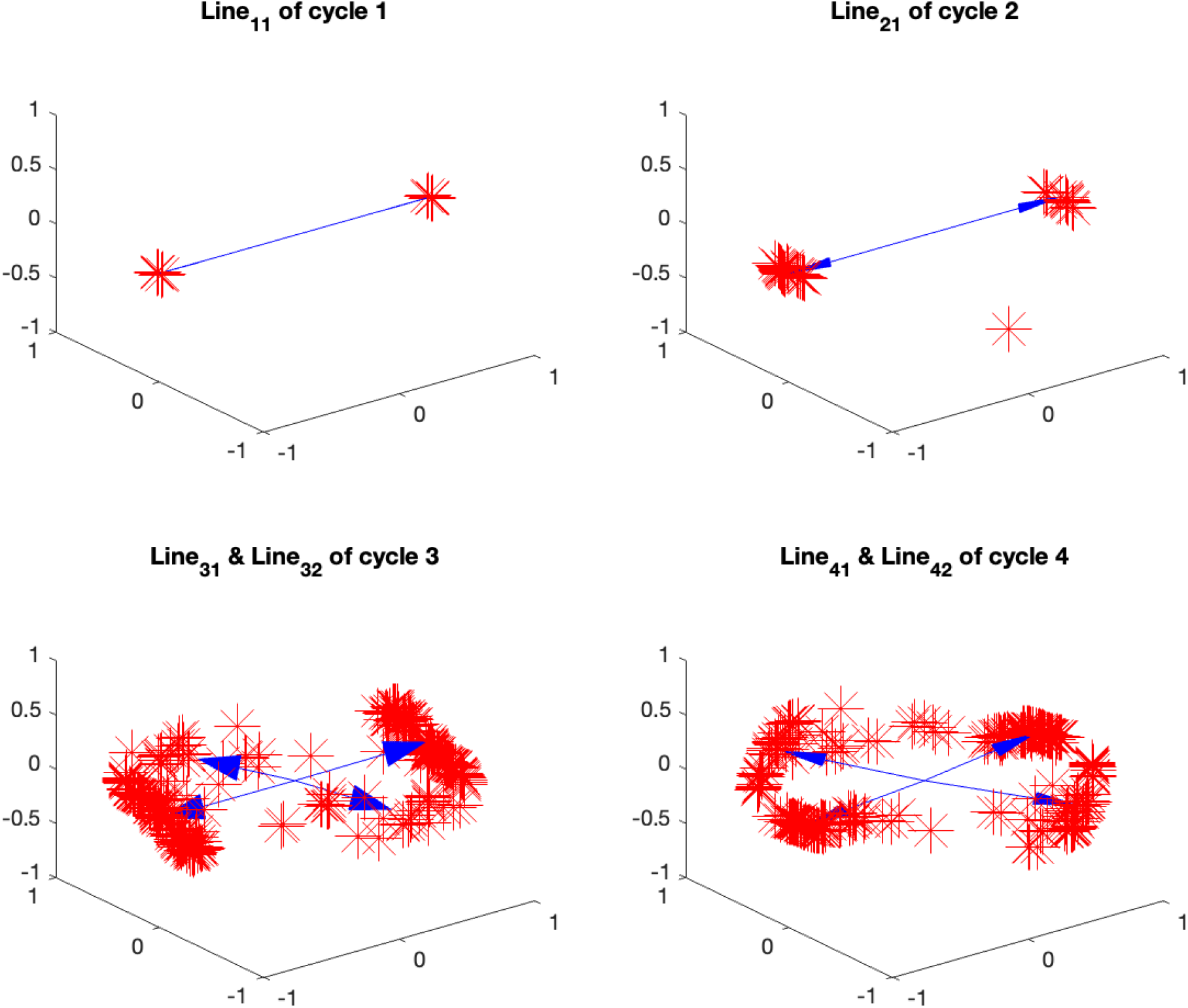
Learned weight vectors forming multiple AIME components with synthetic data after each cycle. In contrast to the Iris data, the synthetic dataset exhibits a more scattered distribution across certain axes. Unlike the patterns shown in Figure 9, where the vectors align along a single line, the synthetic dataset shows that the learned weight vectors span multiple lines (i.e., AIME components) in cycle 3 and cycle 4. The plot for cycle 4 is generated using the masked input data projected onto *line*_31_ from cycle 3 (the counterparts after projecting onto *line*_32_ are omitted in this plot but the idea is similar). These lines are determined with K-Means clustering with *K* = 4 upon subjective observation. This divergence in vector alignment reflects the synthetic data’s diverse and nuanced principal components, where the underlying features do not align neatly along a single axis.

### Numerical outcomes

Tables 2 and 3 show the correlations between the AIME components and true PCs, and the explained variances by the AIME components.

**Table 2:**
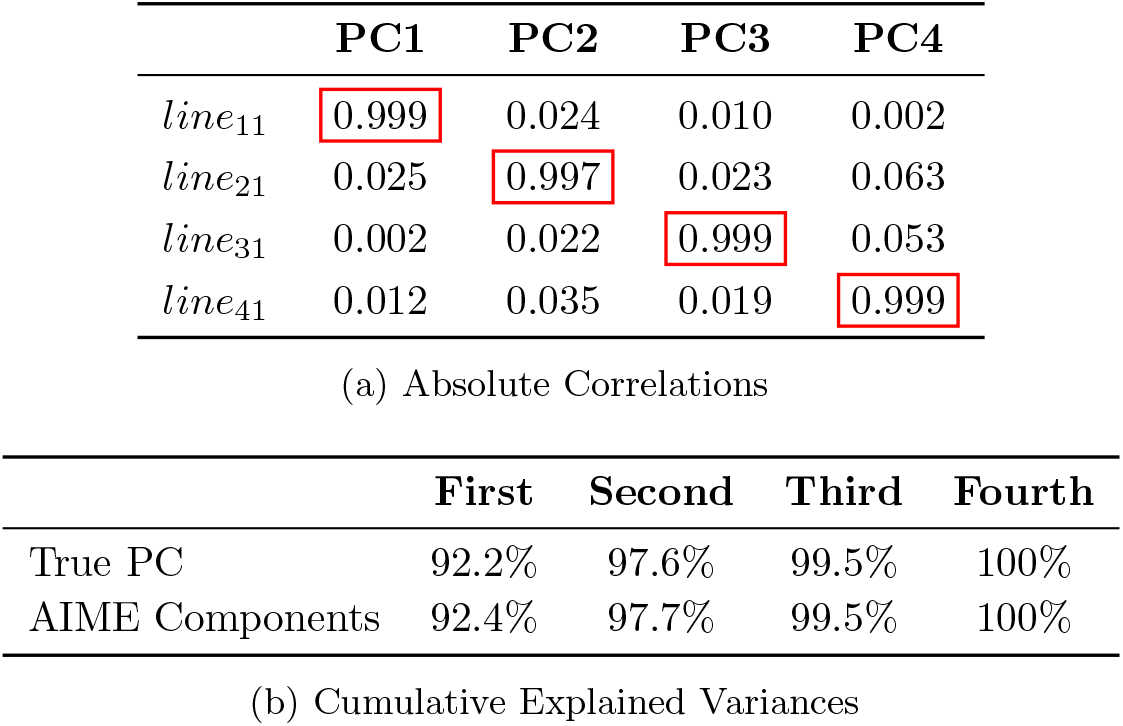
Numerical outcomes of AIME with the Iris data after 4 training cycles. **(a)** shows the cosine similarities between the AIME components and the four true principal components. It is noticeable that the diagonal contains the highest values in the row (circled in red), and all of them are approximately equal to 1, indicating a solid collineation. In **(b)**, the similarly explained variances for both the true principal components and the AIME components exhibit a close alignment, which suggests that along the vectors determined by either method, the data demonstrates nearly identical dispersion. The validation of both measurements ensures the feasibility of AIME.

**Table 3:**
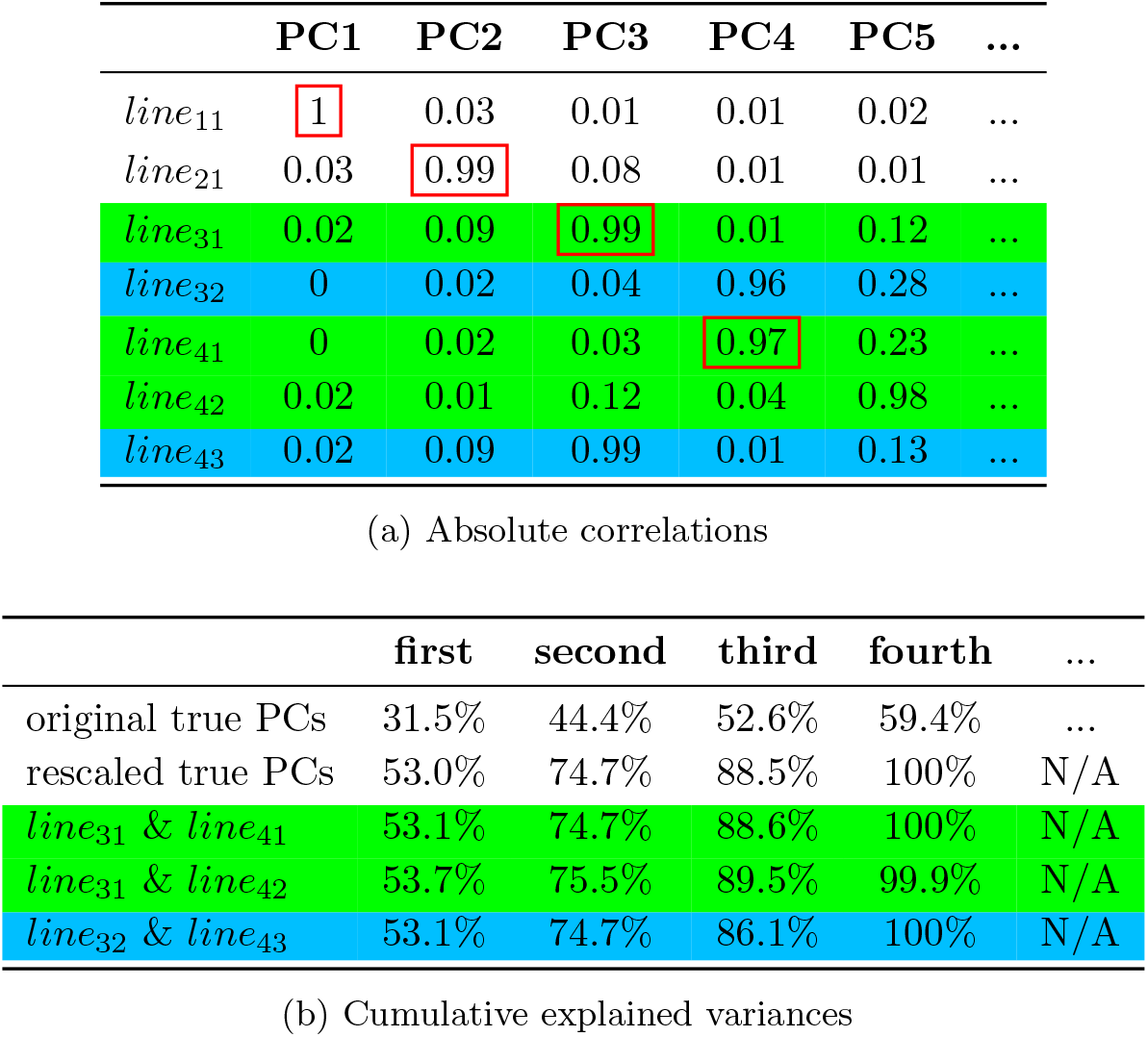
Numerical outcomes of AIME with the synthetic data after 4 learning cycles, where 4 out of 50 principal components are computed. The explained variances of the data provide a valid basis for elucidating the non-uniqueness of the resultant AIME components. The hierarchical relationship between the lines are explained in the diagram 8. From the absolute correlations shown in **(a)**, we see that in cycle 3 and cycle 4, one AIME component well aligns with their corresponding true principal components (i.e., *line*_31_ and *line*_41_, circled in red), while others incline to another directions deviating from the leading PC (i.e., *line*_32_, *line*_42_, and *line*_43_). In **(b)**, we compare the accumulative explained variances between the full set of AIME components and the first 4 true PCs (rescaled for better comparison). The rows highlighted in green indicate AIME components of {*line*_11_, *line*_21_, *line*_31_, *line*_41_} and {*line*_11_, *line*_21_, *line*_31_, *line*_42_} in order, while the blue row is {*line*_11_, *line*_21_, *line*_32_, *line*_43_}. Due to a more uniform dispersion of the synthetic data across neighboring principal components, the resultant AIME components can potentially align with either the present leading PC or the next PC. Under such a circumstance where data is less structured, regardless of the specific orientations of the lines, as long as its direction can explain variances comparable to those of the true principal components, we consider the simulation completely successful.

## 7 Related work

Our primary findings are these: specific biological circuit characteristics cause them to emergently carry out specific recognizable algorithms: hierarchical clustering and component-based feature extraction. We formalized these emergent algorithms and found that they are not closely related mathematically, and yet they derive from a single biological construct. Thus a deep connection between two disparate algorithms has been identified via biological analyses. Other studies in the machine learning and computational neuroscience literatures have identified portions of these findings.

### 7.1 PCA with competitive learning

Our AIME algorithm is based on the modified competitive learning rules, allowing it to extract meaningful components. The concept of using competitive learning for PCA has been explored in previous research involving self-organizing maps (SOM), initially proposed by Kohonen (Kohonen, T., 1982) for categorization.

Ritter et al. (1992) extended Konohen’s traditional SOM to compute *principal curves and surfaces* of the input data, known as nonlinear generalizations of PCA. For ellipsoidal data, principal components emerge as a special case of principal curves, and through competitive learning, their algorithm generates smooth curves or surfaces passing through the center of the data distribution. The learning process involves training the winning weight vectors toward the center of gravity of the input regions.

Kohonen (1995, 1996) then proposed an *Adaptive Subspace Self-Organizing Map (ASSOM)* as a possible alternative to PCA. ASSOM adapts through competitive learning and creates a set of feature representations (i.e., the subspaces) according to different temporal events. In the ASSOM model, the “winning” neural subspace is determined for each timing episode (a set of sampled input vectors drawn from temporal windows) and is updated for each input sample within the episode. The output of each neuron in ASSOM represents the degree of matching between the input data and the subspace learned by that neuron.

In 2014, López-Rubio et al. (2014) presented a novel neural model called *Principal Components Analysis with Competitive Learning (PCACL)*, which improves traditional “local PCA” methods. This approach enables fast execution while preserving the dimensionality reduction capabilities of PCA. During the learning process, the winning neuron is selected based on the projection errors between the input data and the leading eigenvectors of the neural covariances. PCACL explicitly drives neutral weights towards local means and covariance matrices.

### 7.5 Synergistic connections between PCA and clustering

Over the past two decades, several studies have made attempts to generalize clustering and PCA, either from a mathematical standpoint or by incorporating PCA as a key step in clustering, or vice versa. In 2004, Ding and He (2004) made a significant contribution to understanding the mathematical relationship between PCA and K-means clustering, revealing that the PCA dimension reduction automatically performs clustering according to the K-means objective function. Moreover, they proposed that the cluster centroid subspace is spanned by the first *K* − 1 principal directions, indicating that PCA dimension reduction inherently identifies this subspace. Dayal (2009) used axes perpendicular to the leading principal component (PC) to split the data into two halves, which was then used to proceed with hierarchical clustering. Tajunisha and Saravanan (2010) proposed a novel K-Means clustering method by applying PCA in the cluster initialization step. In their algorithm, PCA is solely used to find the initial clusters.

More recently, Mukherjee and Zhang (2022) emphasized PCA’s standalone significance in clustering, demonstrating that it enhances accuracy by reducing intra-cluster distances. They suggested that PCA itself is already a significant advancement in clustering, regardless of the associated (previously, subsequently, or concurrently used) clustering algorithms.

These research studies have extensively explored the direct synergy between PCA and clustering, and significantly enriched their shared mathematical foundations. Our introduction of the AIME algorithm takes a substantial further step, providing a completely novel approach that generalizes these two separate algorithms (components and clustering) to a single underlying algorithm, derived directly from biological circuit structure and operation.

## 8 Conclusions and discussion

We introduced the LI-HC and AIME algorithms, identifying distinct facets of an underlying homogeneous neural architecture. We showed that methods for the statistical operations of clustering, and of components analysis, shared a deep connection, arising as special cases from a single underlying architecture, which was directly derived from biology.

Biologically, the operations of clustering and PCA-like feature extractions through thalamocortical circuits correspond to different characterizations of L-FLIC. In sum, they are:

- Similar in anatomy: both systems share the same local circuits, known as L-FLIC, and the same global circuits, which are the thalamocortical circuits.
- Different in physiology: one key difference lies in the strength of local lateral inhibition, where one system exhibits strong lateral inhibition while the other shows inhibited or weaker lateral inhibition.

Computationally, the algorithms of LI-HC and AIME share the same learning rule but suggest different implementations. The detailed similarities and differences between these two algorithms are illustrated below:

- Similarities: firstly, both the LI-HC and AIME algorithms are rooted under the same learning rule, with a minimal variation. Since competitive learning is a special type of fixed-point iteration, both algorithms involve the convergence to mean vectors of different subsets of input data. Secondly, the neurons’ representations can be the same set of weight vectors. This aligns with the biological reality that both algorithms stem from the same neural architecture. Lastly, both algorithms undergo a process of “input masking” for hierarchical computations. This reflects long-lasting inhibitory processes acting on the core thalamic nuclei.
- Differences: the LI-HC and AIME algorithms exhibit distinct learning processes. One fundamental difference lies in their competitive mechanisms: LI-HC relies on a winner-take-all effect, where weight vectors compete to be selected by subsets of input data. However, AIME operates without such competition, allowing each weight vector independent access to the entire dataset. Additionally, their training processes diverge: LI-HC focuses on the lengths and precise directions of input data, while AIME extends this with a bidirectional spanning of data. Furthermore, their implementations vary. For instance, normalizing the input data to a unit circle is mandatory in LI-HC, whereas AIME does not allow such normalization.

Future directions of AIME may include:

- The current algorithm is well suited to extract meaningful features for linear data, but faces challenges in identifying non-linear manifolds.
- With repeated data simulations, we surprisingly find that there are at most two AIME components found in each learning cycle. This unexpected finding is a topic for ongoing further research.
- A more precise clustering method than K-means may be necessary to accurately identify the AIME components from the learned weight vectors. To avoid the subjective selection of *K*, the clustering module of LI-HC offers a viable alternative for automatically determining their aggregated groups. We have implemented this approach in our repository but do not discuss the details in this paper.
- The current analytic treatment includes both formal proofs and data-driven empirical studies. As a future direction, further research could refine these analyses by incorporating additional theoretical insights and expanding empirical validation across diverse datasets.

## Footnotes

1 The code can be found in the project’s GitHub repository. [Click here for details](https://github.com/charlesl3/LFLIC.git).

2 The original result was in 3D. In Figure 2, we rotated the plots to improve the presentation of the centroids’ relative positions.

